# AVENGERS: Analysis of Variant Effects using Next Generation sequencing to Enhance *BRCA2* Stratification

**DOI:** 10.1101/2023.12.14.571713

**Authors:** Sounak Sahu, Melissa Galloux, Eileen Southon, Dylan Caylor, Teresa Sullivan, Matteo Arnaudi, Josephine Geh, Raj Chari, Elena Papaleo, Shyam K. Sharan

**Affiliations:** Mouse Cancer Genetics Program, Center for Cancer Research, National Cancer Institute, Frederick, MD 21702, USA; Independent bioinformatician, Marseille, France; Cancer Systems Biology, Section for Bioinformatics, Department of Health and Technology, Technical University of Denmark, 2800, Lyngby, Denmark; Cancer Structural Biology, Danish Cancer Institute, 2100, Copenhagen, Denmark; Genome Modification Core, Laboratory Animal Sciences Program, National Cancer Institute, Frederick, MD 21702, USA

**Keywords:** Embryonic Stem Cells, *BRCA2*, Saturation genome editing, Variants of Uncertain Significance (VUS), Multiplexed Assays for Variant Effect (MAVE), Free energy calculations

## Abstract

Accurate interpretation of genetic variation is a critical step towards realizing the potential of precision medicine. Sequencing-based genetic tests have uncovered a vast array of *BRCA2* sequence variants. Due to limited clinical, familial and/or epidemiological data, thousands of variants are considered to be variants of uncertain significance (VUS). To determine the functional impact of VUSs, here we develop AVENGERS: Analysis of Variant Effects using NGs to Enhance BRCA2 Stratification, utilizing CRISPR-Cas9-based saturation genome editing (SGE) in a humanized-mouse embryonic stem cell line. We have categorized nearly all possible missense single nucleotide variants (SNVs) encompassing the C-terminal DNA binding domain of *BRCA2.* We have generated the function scores for 6270 SNVs, covering 95.5% of possible SNVs in exons 15-26 spanning residues 2479-3216, including 1069 unique missense VUS, with 81% functional and 14% found to be nonfunctional. Our classification aligns strongly with pathogenicity data from ClinVar, orthogonal functional assays and computational meta predictors. Our statistical classifier exhibits 92.2% sensitivity and 96% specificity in distinguishing clinically benign and pathogenic variants recorded in ClinVar. Furthermore, we offer proactive evidence for 617 SNVs being non-functional and 3396 SNVs being functional demonstrated by impact on cell growth and response to DNA damaging drugs like cisplatin and olaparib. This classification serves as a valuable resource for interpreting unidentified variants in the population and for physicians and genetic counselors assessing *BRCA2* VUSs in patients.

## Introduction

An array of *BRCA2* sequence variants have been identified due to the availability of sequencing-based genetic tests^1^. Assessing pathogenicity by linking genotype to phenotype is challenging as many of the variants of uncertain significance (VUS) are rare and may only be identified in a single person or family^2–5^. Hence, multiplexed assay for variant effect (MAVE) are necessary to assess a large number of variants experimentally^6–9^. The advent of CRISPR-based SGE has facilitated the creation of programmed variants enabling high-throughput variant screening and multiplexing using Next Generation Sequencing (NGS)^10,11^. Albeit with notable trade-offs, recent CRISPR tools like homology-directed repair^10,11^, base editing^12–14^ and prime editing^15^ have also emerged to classify genetic variants of many cancer-predisposing genes.

Hereditary breast and ovarian cancer (HBOC) results predominantly due to mutations in *BRCA1* and *BRCA2* genes. The exponential increase in sequencing of breast cancer patients as well as individuals at risk of developing the disease has unveiled numerous variants in cancer pre-disposing genes (**Figure 1a**), with several types of mutations identified in *BRCA2* (**Figure 1b**). The clinical variant database, ClinVar, has reported a total of 13,201 *BRCA2* SNVs to date, with 59% still considered to be VUS^16^ (Clinvar update: 30 Oct, 2023) (**Figure 1c**). As the number of unique variants observed is expected to increase with improved sampling in the coming years, it is crucial to develop more efficient methods of variant assessment and classification. Functional assays have been developed to study the effect of *BRCA2* VUSs on homologous recombination, proliferation, and sensitivity to chemotherapeutic drugs, resulting in an impressive collection of data^17–22^. Despite a concerted and interdisciplinary community effort, the classification of less than a few thousand variants has been achieved to date. Only a handful of these variants have been studied using genetic murine models, which are time consuming and resource-intensive^23–25^.

**Figure 1:**
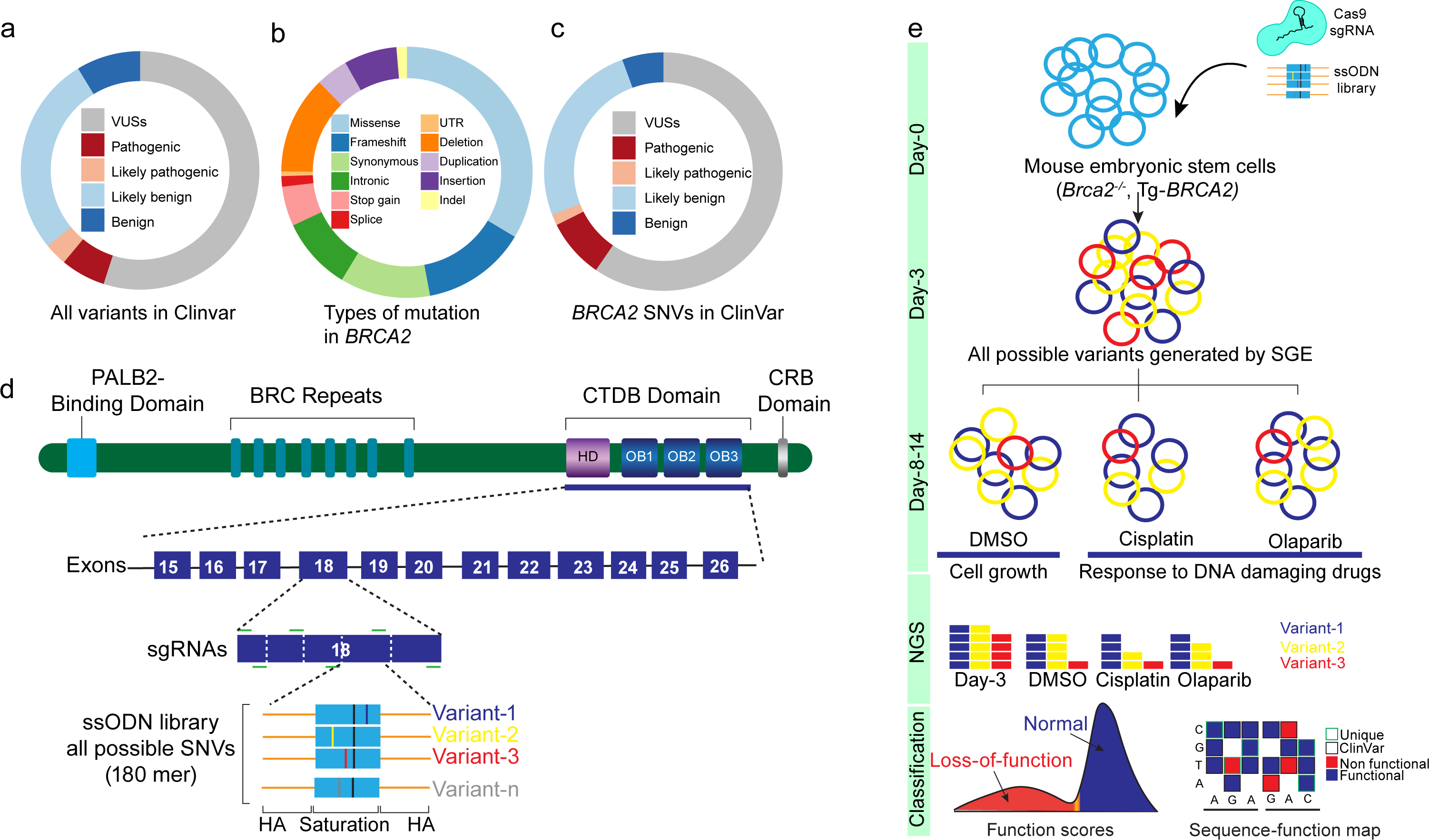
Reducing uncertainty in *BRCA2* variants of uncertain significance (VUSs) using Saturation Genome Editing (SGE) Donut plots showing the distribution of **(a)** different classes of genetic variants identified for cancer predisposition genes and reported in ClinVar, the Clinical Variant database (n = 2,226,284). 55% of all variants are still considered to be VUS. **(b)** Different types of mutation identified for *BRCA2* (n = 23060); 33.5% missense variants are identified. **(c)** Distribution of *BRCA2* SNVs identified in ClinVar (n=7863 VUS, 1064 Pathogenic, 197 likely pathogenic,3339 likely benign and 738 benign). 59% of *BRCA2* SNVs is still considered to be VUS. **(d)** Schematic representation of the strategy for sgRNA and oligo donor design to saturate the entire CTDB domain of *BRCA2* encoded by exons 15-26. Depending on the availability of sgRNAs (green lines), we used oligo pools as repair template to saturate 12 exons across the entire CTDB domain. Each oligo-pool comprises of 180-mer long ssODNs containing a synonymous mutation at the PAM site (black line) and degenerate nucleotides (blue, yellow and red lines) containing all three non-wildtype nucleotides per position across the region of saturation. (HA=Homology arm represented with an orange line, CRB = C-terminal RAD51-binding domain). **(e)** Schematic representation of the experimental workflow of CRISPR-SGE experiments where mouse ES cell line expressing a single copy of human *BRCA2* (*Brca2*^−/−^,^Tg[*BRCA2*])^) were transfected with sgRNA-Cas9 plasmids, and successful transfection was confirmed by GFP fluorescence. Two groups of cells were then analyzed: one on day 3 to determine initial variant representation and another on day 14 days with or without treatment of DNA damaging agents (cisplatin and olaparib) to evaluate the variant abundance. Variants that decreased in abundance over time are considered non-functional, while those that did not significantly deplete are deemed functional.

BRCA2 is a 3418-amino acid protein consisting of several functional domains such as the N-terminal domain^26,27^ that interacts with Partner and localizer of BRCA2 (PALB2), eight BRC repeats in the middle of the protein and a C-terminal DNA binding (CTDB) domain consisting of a helical domain (HD), tower domain and three oligosaccharides binding (OB) domains (**Figure 1d**). The BRC repeats as well as the C-terminal RAD51-binding (CRB) domain of *BRCA2* can bind to RAD51, which is required for repair of DNA double strand breaks by homologous recombination (HR)^28–32^. Because of its critical function in maintaining the genomic integrity, BRCA2 is essential for cell proliferation and viability of mouse embryonic stem cells (mESCs)^10,33,34^. Thus, assessing fitness defect provides a reliable parameter for identifying loss-of-function (LoF) variants. We have stably integrated human *BRCA2* into the mouse genome that does not contain *Brca2.* This enables us to engineer a single variant per cell and investigate the effect of these variants on regulatory activity, splicing and protein function^35^.

Using AVENGERS, we have classified all possible SNVs in *BRCA2* exons 15–26 encoding the CTDB domain spanning residues 2479–3216 due to the predominance of known ClinVar-reported pathogenic missense variants in this domain ^36,37^. For SGE, we used a mouse ES cell line expressing a single copy of human *BRCA2* (*Brca2*^−/−^,^Tg[*BRCA2*]^)^35^ along with either a single or a combination of guide-RNAs (sgRNAs) and spCas9 along with a library of 180 bp long single-stranded oligo donors (ssODN) for variant generation using homology-directed repair (HDR) (**Figure 1d**). Pools of cells were sampled at an initial time point (day 3) and a later timepoint (day 14) either without any drug treatment (DMSO only) or after treatment with cisplatin (damages DNA by forming DNA adducts) or olaparib (poly (ADP-ribose) polymerase inhibitors, PARPi) treatment. We calculated the relative frequency of variants based on their NGS read counts in the pool of cells. Classifying genetic variants based on PARP inhibitor sensitivity, which reflects a defect in HR, could also provide valuable, patient-specific clinical insights and actionable treatment options^38^. The dropout frequency of SNVs is defined as function scores which were analyzed using a Gaussian-mixture-model to calculate their probability of impact on function (PIF) (**Figure 1e**). Following The American College of Medical Genetics and Genomics (ACMG)/Association for Molecular Pathology (AMP) sequence variant interpretation guidelines^39,40^, we derived the strength of evidence that could be applied for BRCA2 clinical variant interpretation based on the performance of our classified variants with known clinical relevance.

## Results and Discussion

### SGE coupled with cell viability and chemotherapeutic drug response measure BRCA2 function

Depending on the availability of sgRNAs, we used 48 different oligo pools as HDR template for SGE of 12 exons and ∼3-7bp of adjacent intronic sequences that encode the entire CTDB domain. Each oligo-pool is comprised of 180-mer long ssODNs (ranging from 87-339 ssODNs per pool) that contain: a) a synonymous mutation at the PAM site to block recutting by Cas9, which also acts as a fixed HDR marker during sequencing analysis and b) degenerate nucleotides containing all three non-wildtype nucleotides per position along the 30-50 bp region of saturation (**Figure 1d**). Transfections for SGE experiments were performed on day 0 and genomic DNA were isolated for amplicon sequencing at days 3 and 14 to measure the frequency of variants present in each pool^35,41^ (**Figure 1e**). Using this approach, we have recovered 95.5% of all possible missense variants (6270 out of 6544 SNVs) **(Supplementary figure 1a).** The rate of indels varied within the range of 5% to 45% of the total read count, while HDR rates ranged from 2% to 16% across different exons depending on the respective sgRNAs **(Supplementary Figure 1b-c)**. Notably, we observed a robust correlation in read counts for SNVs generated by HDR from two independent replicates at day 3 **(Supplementary Figure 1d)**. This correlation enabled us to use initial read count frequency in our calculations and compute relative dropout frequency, which we have termed "function scores (FS)".

We have assigned function scores (FS) to 6270 variants by taking the log2 ratio of each SNV’s frequency on day 14 using the day 3 samples as a baseline for fold change calculation. The positional biases in editing rates were corrected and the subsequent FS values were normalized between oligo pools and across different exons such that the median synonymous SNV and nonsense SNV matched the global medians. Additionally, FS values were computed by considering dropout frequencies following treatments with cisplatin and olaparib at day 14. Our rationale for incorporating the response to DNA damaging agents is twofold: to introduce an additional parameter for robustly categorizing *BRCA2* variants and to refine SNV filtering to prevent erroneous functional classification. Our working hypothesis is that variants displaying normal cellular fitness will survive in the presence of these drugs, while loss-of-function (LoF) variants will not.

We found a strong correlation between FS values derived from cellular fitness and those from the response to cisplatin and olaparib (ρ= 0.84 for DMSO vs Cisplatin, ρ= 0.85 for DMSO vs olaparib, and ρ= 0.87 for olaparib vs cisplatin) **(Supplementary Figure 2a)**. FS values were bimodally distributed, and we employed a two-component Gaussian Mixture Model (GMM) to calculate the probability of impact on function (PIF).

The PIF scores were used to classify the 6270 SNVs based on the cell fitness data that survive in the pool at day 14. Similarly, we generated integrated FS values by taking a weighted-mean of FS values from cell fitness (DMSO-treated) and drug response (cisplatin-treated and olaparib-treated) data. By employing GMM, we categorized variants based on the integrated values, revealing that 5968 (95.1%) SNVs exhibited strong concordance, while only 302 (4.8%) SNVs showed an opposite classification with cell fitness data. Consequently, we filtered out 302 SNVs as "uncertain" that could not confidently be scored **(Supplementary Figure 2b-d)**. In summary, our approach, which integrates assessments of cellular fitness and responses to DNA damaging drugs, allows for the accurate classification of BRCA2 variants.

### Unraveling the pathogenic spectrum of *BRCA2* variants

We observed that the majority of frame-shifting indels in each of our experimental pools exhibit a significant impact on cellular fitness, emphasizing the efficacy of our approach in accurately categorizing LoF variants **(Supplementary Figure 3a)**. FS values for all nonsense SNVs scored below −1.11 (N = 310, median = −2.00), whereas synonymous SNVs scored above −0.95 (N = 1266, median = −0.02) (**Figure 2a**). The posterior probability computed from GMM were used to categorize SNVs by setting thresholds for PIF scores as follows: *P*_nf_ > 0.95 = ‘non-functional’, 0.01< *P*_nf_ < 0.95 = ‘indeterminate’, *P*_nf_ <0.01 = ‘functional’ **(Supplementary Figure 4a)**. The FS values for all variants strongly correlate between the replicates clearly demarcating the functional and non-functional SNVs **(Supplementary Figure 4b)**. We also observed a clear separation between synonymous and non-sense SNVs across all the exons, which strengthens the accuracy of our functional categorization **(Supplementary Figure 4c)**.

**Figure 2:**
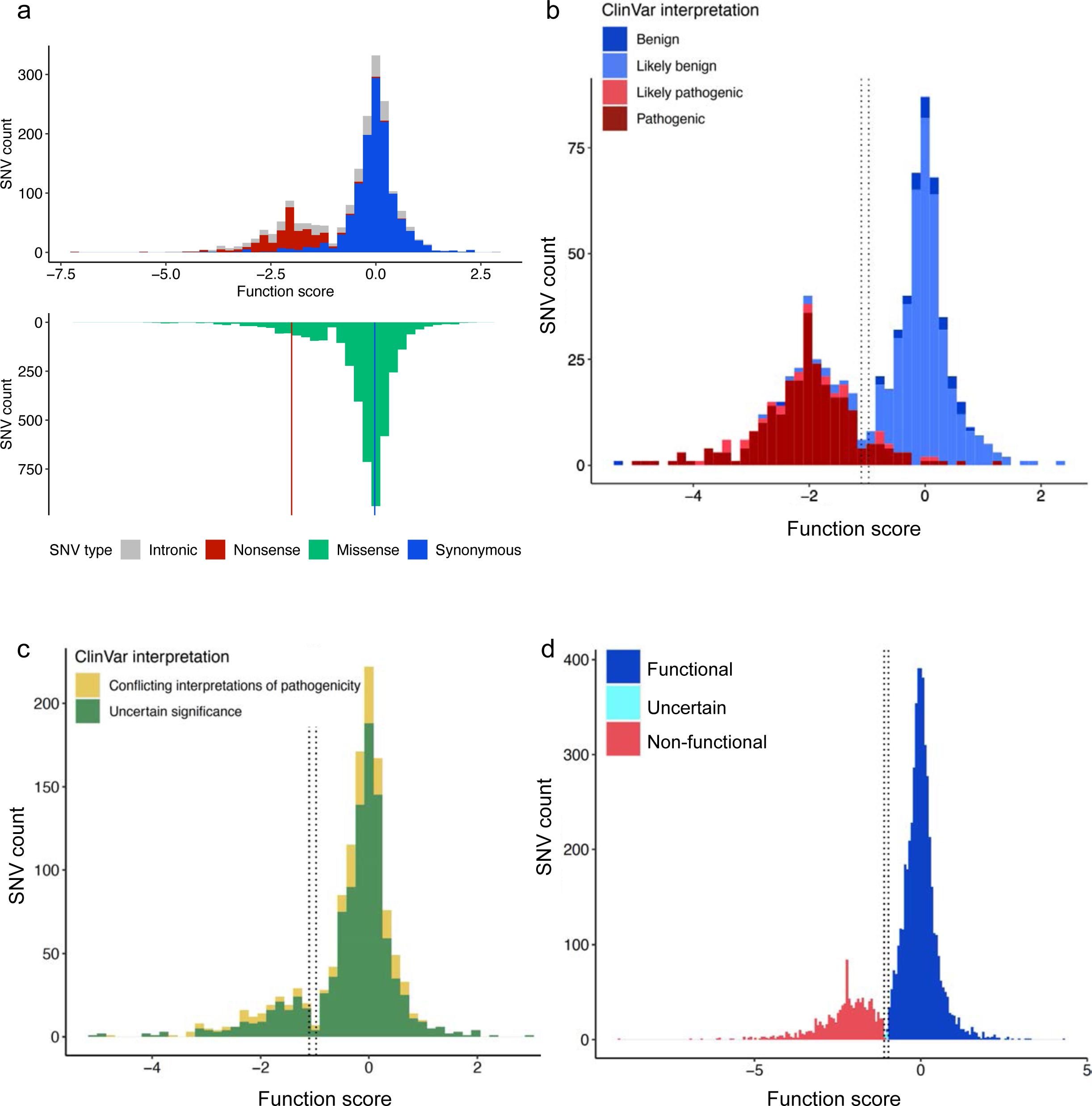
AVENGERS accurately determine clinical interpretations of BRCA2 SNVs. **(a)** The distribution of SNV function scores for Nonsense (red), Synonymous (blue), intronic (grey) and Missense (light green). The dashed line represents the median of the nonsense SNVs (red) and synonymous SNVs (blue). **(b)** Histogram showing distribution of SNV function scores categorized by ClinVar interpretation (n = 717), with at least a ‘1-Star’ review status in ClinVar and either a ‘Pathogenic’ or ‘Benign’ Interpretation (including ‘Likely benign and likely pathogenic). The dashed lines represent functional and non-functional classification thresholds derived from the Mixture Modeling. **(c)** Histogram Illustrating the distribution of 1069 SNVs classified as Variants of Uncertain Significance (VUS) alongside 233 SNVs with Conflicting Interpretations of Pathogenicity in ClinVar. The bimodal distribution indicates our capacity to categorize them as either functional or non-functional, demonstrated by the dashed lines derived from Mixture Modeling. **(d)** Distribution of function scores for all possible SNVs (n = 5968) across exon 15-26 from the c-terminal DNA binding domain. We categorized 4881 SNVs to be functional (81.7%) and 1087 SNVs to be non-functional (18.2%).

We have classified 1165 synonymous SNVs (92%) as functional and 283 nonsense SNVs (91.2%) as non-functional (**Supplementary Figure 4d)**. We hypothesize that potential impact on splicing may explain the discordance in the interpretation of a small percentage of variants, specifically where synonymous SNVs behave as non-functional and protein-truncating variants behave as functional. For example, synonymous SNV p.Pro3039Pro (c.9117G>C and c.9117G>A) at the canonical splice site leads to premature protein truncation caused by exon 23 skipping^42^, which explain the non-functional class of this synonymous SNV. Similarly, we have previously identified p.Trp194X(c.809G>A) mutation, predicted to create an N-terminal truncation, produces a fully functional BRCA2^43^. This highlights that, despite predictions of premature protein truncation, some alleles may express alternatively spliced transcripts that result in functional proteins.

We next investigated the ClinVar variants that are expert-curated for BRCA2 which could be a valuable resource for verifying the accuracy of our classification. Out of the 301 SNVs classified as ‘pathogenic’ or ‘likely pathogenic’ in ClinVar, which are also included in our classifications, 255 (84.7%) were categorized as ‘non-functional’, 31 as ‘functional’ and the remaining 15 as ‘uncertain’. Conversely, among the 444 SNVs labeled as ‘benign’ or ‘likely benign’ in ClinVar, 412 (92.79%) SNVs were classified as ‘functional’, 19 as ‘non-functional’ and 13 as ‘intermediate’ (**Figure 2b, Supplementary Figure 4e)**. Additionally, we provided interpretation to 1069 VUSs with identification of 865 (80.92%) as ‘functional’ and 151 (14.12%) as ‘non-functional’ (**Figure 2c**). Furthermore, we categorize 233 SNVs with ‘conflicting interpretations of pathogenicity’ as 177 (75.96%) ‘functional’ and 45 (19.31%) ‘non-functional’ (**Figure 2c**). Using this approach, we classified 4881 SNVs to be functional (81.7%) and 1087 SNVs to be non-functional (18.2%) across 12 exons from the CTDB domain, (**Figure 2d, Supplementary Figure 4f)**. We observed an AUC value of 0.96 with 92.2% sensitivity and 96% specificity of our model to accurately classify pathogenic/likely pathogenic and benign/likely benign variants annotated in ClinVar. Similarly, AUC value of 0.97 was observed in classifying synonymous and nonsense SNVs with 93.5% sensitivity and 95.2% specificity **(Supplementary Figure 5a)**. We have also compared FS values of ClinVar variants to commonly used computational predictors like CADD^44^, BayesDel^45^, REVEL^46^, PRIOR, EVE^47^ and AlphaMissense^48^. While there is generally a broad consensus among these models in predicting LoF variants, some functional SNVs were often interpreted as pathogenic in computational predictors. We observed AUC ranging from 0.71 to 0.92, outperforming predictions from computational models and functional assay results **(Supplementary Figure 5 a-b)**. In summary, the strong concordance of our results with ClinVar helps ensure reliability of variant classification for clinical decision-making and genetic counseling. The AVENGERS map revealed the pathogenicity spectrum of all possible SNVs across the CTDB domain and splice site variants (**Figure 3**).

**Figure 3:**
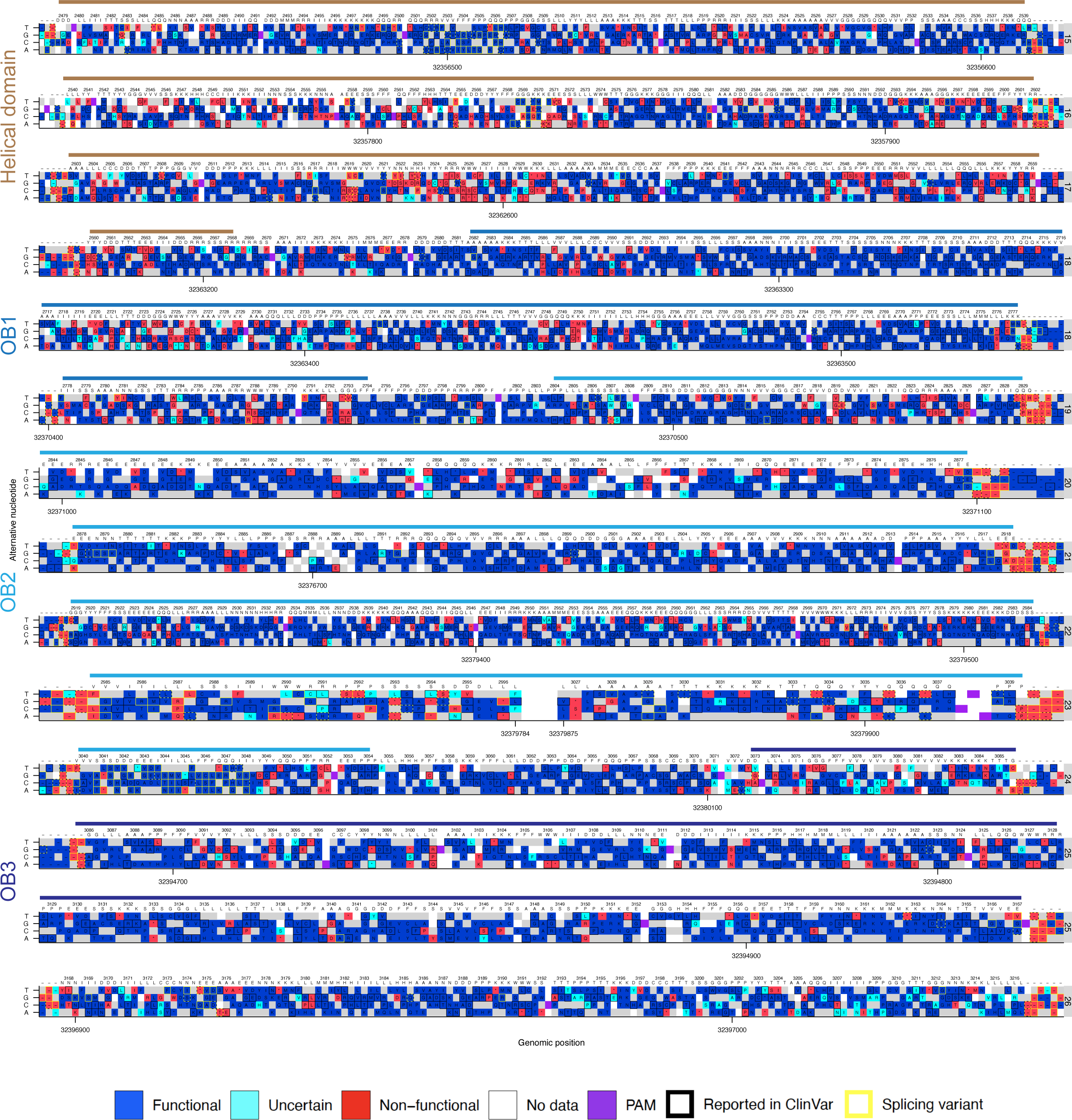
AVENGERS map for 6270 *BRCA2* SNVs across 12 exons spanning the C-terminal DNA binding (CTDB) domain. The Sequence-function map unveils the pathogenic range of all potential SNVs in the BRCA2 CTDB Domain. The box color signifies Functional (Blue), Non-Functional (Red), and Uncertain (Turquoise Blue) class. The wild-type nucleotide is shown in Gray, while white boxes indicate excluded nucleotides. A black border surrounding the box denotes SNVs reported in ClinVar, and a dashed border represents SNVs predicted to affect splicing with SpliceAI Score > 0.2^61^. The alphabet denotes the single-letter amino acid code and their change due to SNVs. The numbers at the top denote the amino acid residue and the bottom indicate the nucleotide position on chromosome 13 as per the Genome Reference Consortium (Human GRCh38). The solid line depicts individual functional domains. Residues 2830-2843 (exon20) and 2997-3026 (exon 23) were excluded due to lack of efficient sgRNAs for SGE.

### Structural interpretation aligns with *BRCA2* variant pathogenicity

Furthermore, we applied a framework for variant interpretation based on in-silico structural methods, i.e., MAVISp (Multi-layered Assessment of Variants by Structure for proteins)^49^ to investigate the effects of the *BRCA2* variants in the CTDB domain and that are reported in COSMIC^50^, cBioPortal^51^, and ClinVar^36^. MAVISp allows to scrutinize variants’ effect using not only pathogenicity scores as EVE^47^ and AlphaMissense^48^, but also to annotate the impact of variants on different protein features, such as structural stability, and protein-protein interactions. CTDB domain interacts with *Deletion of Split hand/split foot* 1 (DSS1/SEM1) which is essential for the stability of BRCA2^27^. We focused on the effects on structural stability or on the interaction between BRCA2 and DSS1/SEM1. We observed a strong concordance between MAVISp-based saturation scans for the CTDB and AVENGERS and Alpha-missense (**Figure 4a**). For 92 of the non-functional variants, we identified an effect due to changes in structural stability according to the Rosetta and FoldX predictions. The 38 non-functional variants the effects seem to be related to changes in the interaction with DSS1/SEM1 (**Figure 4b-e**). Moreover, p.Gly2784Val, p.Ala2786Pro and p.Gly2739Arg in the CTDB domain are predicted to destabilize both stability and the DSS1/SEM1 interaction.

**Figure 4:**
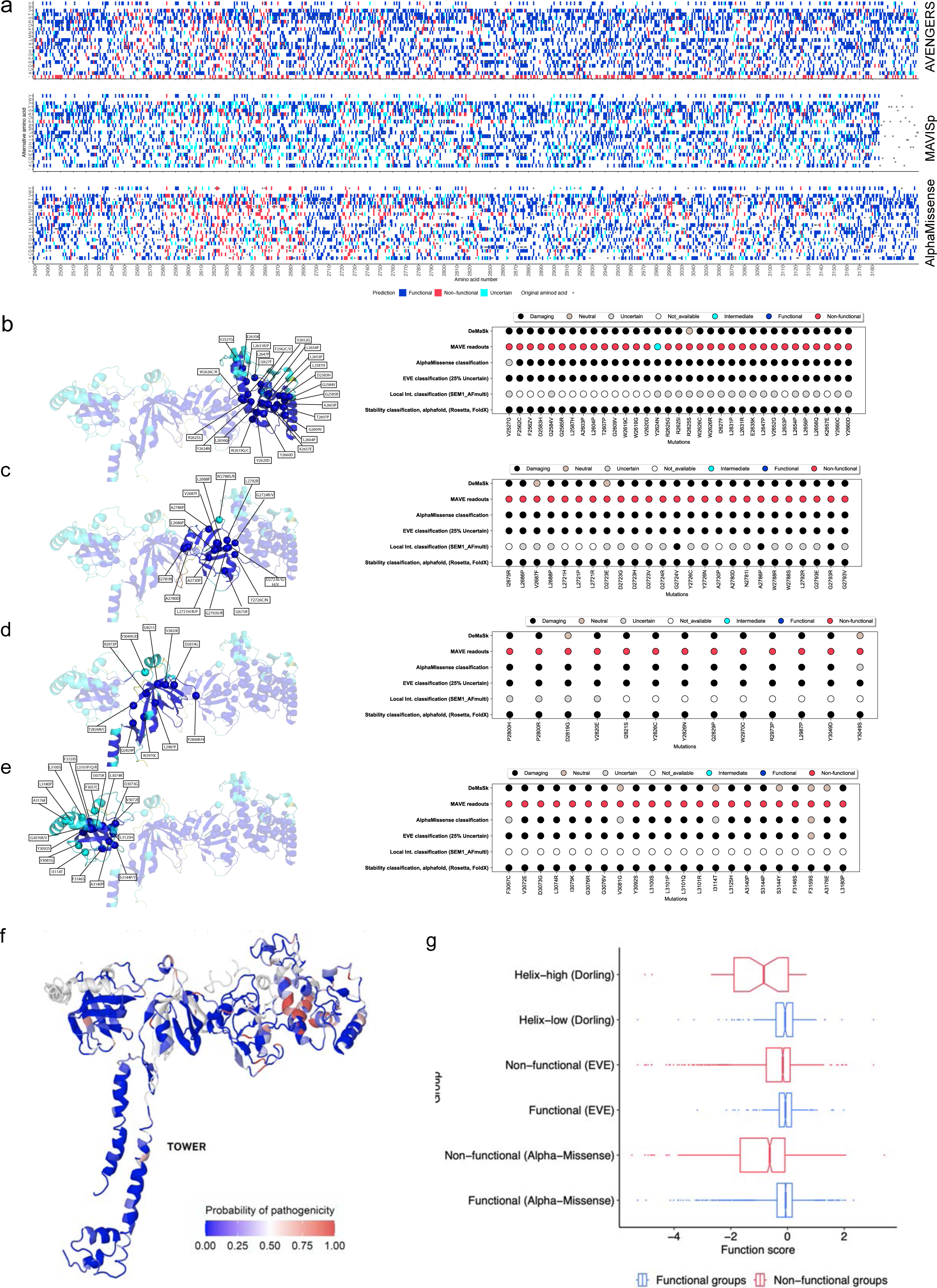
Structural prediction and adaptation of SGE classes for breast cancer risk estimation. MAVISp predictions demonstrate concordance with SNVs classified as non-functional according to MAVE assay and their destabilizing status with DSS1 interaction. **(a)** Sequence function map demonstrating strong concordance between AVENGERS, MAVISp and Alphamissense across residues 2479 to 3186 for all possible amino acid changes. Blue box represent functional/stabilizing, black as non-functional/des-stabilizing and turquoise blue as uncertain. The original amino acid residue is represented as a circle. **(b-d)** The dot plots reveal the consequences of individual SNVs derived from pathogenicity predictors like DeMAsk, EVE, and AlphaMissense. The dot plots also include the effects predicted for each SNV with the stability and local interaction modules of MAVISp. The Structural 3D plots are derived from BRCA2 CTDB domain from Alphafold depicting the helical domain **(b),** OB1 **(c),** OB2 **(d)** and OB3 **(e).** The color code of the 3D structure plot is derived from AlphaFold and correlated with the pLDDT score, a score to provide information about the confidence of the prediction of a specific residue, in particular regions with a pLDDT score <50 are colored in orange, regions with a pLDDT score between 50 and 70 are colored in yellow, regions with a pLDDT score between 70 and 90 are colored in light blue, regions with a pLDDT score > 90 are colored in blue. In MAVISP we used structures or portion of structures with a pLDDT score > 70. **(f)** The probability of pathogenicity of missense variants across the CTDB is displayed on the BRCA2-DSS1 complex structure (PDB: 1MIU). Residues within helical domain are among the position’s intolerant to SNV changes and variants in the tower domain are mostly tolerated. **(g)** Function scores from AVENGERS can predict clinical pathogenicity derived from BRIDGES consortium dataset^3^ and their comparison to AlphaMissense^48^ and EVE^47^ score prediction (n = 36 helix-high, n = 220 helix-low SNVs).

Furthermore, the predictions on variants at the p. Leu2721 residue changed to His, Pro, and Arg, and p.Asp2723 residue changed to Gly, His, and Val in terms of stability are consistent with the non-functional readout of AVENGERS and align with AlphaMissense prediction (**Figure 4 d,e**). This concordance is displayed in the sequence-function map which portrays the effects of changing individual residues within the CTDB domain to all amino acid residues **(Supplementary Figure 6 a-d)**. The p.Leu2721His was previously characterized as deleterious based on HR assay and its sensitivity to PARP inhibition. Taken together, we demonstrate MAVISp predictions for *BRCA2* variants align with AVENGERS, providing a mechanistic explanation underlying the impact of SNVs resulting in non-functional experimental readouts.

We overlaid the non-functional SNVs In the BRCA2 3D structure^27^ and revealed a region spanning from residues 2619 to 2630 within the helical domain that is highly intolerant to missense variation (**Figure 4f**). Notably, the p.His2623 residue clusters within the helical domain, which has been previously identified to contain deleterious variants^14^. Additionally, residues from 2720 to 2726 within the OB1 fold also exhibit intolerance to variant changes. In this region, several pathogenic BRCA2 variants, such as p.Thr2722Arg and p.Asp2723His, have been identified. Interestingly, the OB2 and OB3 regions are more tolerant to missense variation, particularly the Tower domain within the OB2 fold, where no pathogenic variants have been previously reported.

The mixture-models revealed 92% sensitivity and 96% specificity to accurately classify ClinVar variants, and the positive likelihood ratio (LR+) based on ClinVar is 23.64 and the negative likelihood ratio (LR-) is 0.08. Furthermore, to estimate the evidence strength of our assay, we calculated the Odds of Pathogenicity (OddsPath) value of our model, based on the recommendations for application of the functional evidence PS3/BS3 criterion using ACMG variant interpretation framework^39,40^. Our OddsPath value for pathogenic variant is 23.35 equivalent to PS3 level of evidence and for benign variant is 0.006 equivalent to BS3 level of evidence.

To assess whether FSs predict variants driving breast cancer, we correlated our variant classification with a substantial dataset from the Breast Cancer Association Consortium BRIDGES project^2^. This included 59,538 breast cancer cases and 53,165 controls, with 5,467 carriers of 1,425 unique *BRCA2* variants. Notably, we identified 9 SNVs as non-functional in our dataset that were classified as (likely) pathogenic on ClinVar or by ENIGMA expert guidelines. For instance, p.Leu2510Pro (c.7529T>C) was identified in 2 cases and 1 control, and its deleterious effect was previously characterized^35,43,52^. Other SNVs from the helical domain include p.Ile2627Phe(c.7879A>T), p.Leu2647Pro(c.7940T>C), p.Leu2653Pro(c.7958T>C), p.Ile2675Val(c.8023A>G). Other SNVs include p.Asn3124Ile(c.9371A>T) identified in 11 breast cancer cases and 1 control population, p.Gly3076Arg(c.9226G>A) identified in 3 cases and 0 controls, with relative risk estimates similar to truncating variants, confirming their non-functional nature. Correlating breast cancer risk association results for individual SNVs with a frequency between 0.1% and 5% based on population samples in the BRIDGES dataset, we found p.Ala2717Ser(c.8149G>T) (98 cases and 130 controls), p.Val2728Ile(c.8182G>A) (219 cases and 208 controls), p.Glu2856Ala(c.8567A>C) (133 cases and 172 controls), p.Ala2951thr(c.8851G>A) (299 cases and 340 controls) that have a oddsRatio of nearly 1 and are functional in our assay. To summarize the population data, we observed a good concordance to the Helix-High SNVs identified in the BRIDGES dataset to be non-functional and variants with low Helix score are mostly functional in our assay (**Figure 4g**). In summary, our population data demonstrated good concordance with non-functional SNVs in the BRIDGES dataset, suggesting the clinical applicability of our functional assay for patient management in the future.

### AVENGERS scores align with high-throughput orthogonal assays

While our functional categorization has shown strong agreement with ClinVar-classification, it is important to exercise caution when drawing definitive conclusions about the pathogenicity of variants. Reproducible yet unanticipated effects, such as editing at off-target sites, could be a potential caveat in CRISPR-based assays. To address this caveat, we present a holistic approach to compare our SGE data to previously reported orthogonal assays. Several of the sgRNAs used in our SGE experiments were previously used to saturate a few codons of BRCA2 using Prime Editing (PE), known to reduce indel formation and off-target editing^15^. We observed 131 out of 156 SNVs (84%) to be concordant between our AVENGERS classification and PE, of which 116 SNVs were functional and 15 SNVs were nonfunctional **(Supplementary Figure 7a)**. As an independent validation, we used 155 SNVs previously generated using a BAC-recombineering-based method and classified these using our mouse ES cell line^20,33^. We observed 91.6% of the SNV classification matching our SGE dataset, that includes 75 functional and 56 nonfunctional variants concordant between both the datasets (**Supplementary Figure 7b)**. Furthermore, our comparison with 117 SNVs previously classified using the MANO-B assay^21^ based on their response to different PARP inhibitors (olaparib, rucaparib, niraparib) and carboplatin revealed 82% of the SNV classification matches our SGE class. Out of 21 discordant SNVs, we found that one-third of these SNVs fell in the Tower domain of *BRCA2* where SNVs were identified to have sensitivity to PARP inhibitors (fclass-3 or 4). However, SNVs identified in this region had no effect on HR^19^, in agreement with our functional classification **(Supplementary Figure 7c)**. BRCA2 variants associated with defective homologous recombination (HR) render cells sensitive to PARP inhibitors, making them a target for precision cancer therapy. We observed 43 out of 47 SNVs to show concordance to a recently reported BRCA2 variant classification based on the ability to perform HR and integrative in silico prediction^53^. Only two HR-proficient SNVs were classified as non-functional by AVENGERS (p.Gly2593Glu and p.Tyr2601Cys) **(Supplementary Figure 7d)**. In addition, our comparison to a large dataset for several ClinVar variants and their ability to perform HR also showed strong comparison. Interestingly, we observed that SNVs in the Tower domain that can perform HR^19^ are functional in our SGE-based classification **(Supplementary Figure 8a)**.

Imposing a stringent threshold to calculate pathogenicity, we have categorized 302 SNVs as “uncertain” which show discordance between cell fitness-based classification and drug-response. However, these “uncertain” variants that typically survive but exhibit vulnerability to DNA damaging agents could potentially be hypomorphic variants, warranting further investigation **(Supplementary Figure 9a)**. We identified p.Tyr2624Cys (c.7871A>G) to have normal cell fitness but was sensitive to drugs, concordant with reduced HDR^54^ and its pathogenicity identified to be high risk in breast cancer patients^2^. Several BRCA2 SNVs were classified based on their intermediate level of HR activity and associated with moderate risk of breast cancer. This includes p.Gln2925Arg (c.8774A>G) and p.Leu2972Trp (c.8915T>C) which we classified as functional based on integrated dataset, but these are sensitive to either one or both the drugs. The p.Asp2606Gly (c.7817A>G) and p.Gly2812Glu (c.8435G>A) were functional in AVENGERS, despite their intermediate HDR activity. Other hypomorphic variants identified to contribute to a moderate risk of breast cancer^55^ include p.Gly2508Ser and p.Ala2717Ser in the functional class, but p.Tyr3035Ser falls in the indeterminate zone. The p.Gly2609Asp(c.7826G>A) variant located in the coding exon 16 of *BRCA2* had reduced HR activity with a prediction of being likely deleterious^18^ and sensitive to PARPi^21^, concordant with our hypomorphic classification.

In summary, AVENGERS represents a high-throughput MAVE approach to classify missense variants compared to traditional medium and low-throughput assays^22^, offering valuable insights into yet-to-be-identified variants. Despite the inherent limitations of SGE, our approach successfully elucidates the phenotypic effects of 6270 *BRCA2* SNVs spanning the region encoding the CTDB domain. By integrating cellular fitness and response to DNA damaging agents along with structural predictions, we have established that our functional classification aligns closely with ClinVar classification with concordance to other MAVEs and orthogonal functional assays.

## Methods

### 1. Mouse embryonic stem cell culture

Mouse ESC line expressing a single copy of human *BRCA2* (*Brca2*^−/−^,^Tg[*BRCA2*]^) [Clone: F7/F7] ^35^ was cultured on feeder cells (SNLP) that express leukemia inhibitory factor (Lif) and puromycin and neomycin resistance genes^33^. The maintenance medium (M15) consisted of Knockout DMEM (Gibco) supplemented with 15% FBS (HyClone), penicillin-streptomycin-glutamine (Gibco) and 0.1 mM β-mercaptoethanol. The media was changed every day and cells were trypsinized and passaged when ESCs reached 80% confluency.

### 2. Guide RNA design and cloning

20 bp of crRNA sequences targeting exons 15-26 of *BRCA2* were designed for high on-target score using www.benchling.com and synthesized by Integrated DNA Technologies (IDT). The single guide RNAs (sgRNAs) were cloned into PX330 or PX458 (Cas9-P2A-GFP) vectors, that expresses the gRNA from a U6 promoter and Cas9 expression cassette^56^. Complimentary oligonucleotides were annealed, phosphorylated, diluted, and ligated into *Bbs*I-digested plasmid^56^. sgRNAs were selected based on the option to introduce a synonymous variant at the PAM sites. The plasmids were confirmed by Sanger sequencing using U6 primers for the correct integration of crRNA sequences.

### 3. Nucleofection for CRISPR-Cas9 based saturation genome editing

We prepared mESCs for nucleofection by culturing them in a 10 cm culture dish one day before the procedure to ensure they were actively dividing. Subsequently, we performed nucleofection on 3.3×10^6^ mESCs using a combination of 3 µg of plasmid containing the desired sgRNA and Cas9-expressing cassette, along with 6.5 µg of ssODN oligo, following the manufacturer’s instructions for the Lonza nucleofector 2B (Program: A030). To facilitate this process, we synthesized libraries of ssODNs for each sgRNA at a concentration of 50 pmol/oligo. These ssODNs were precisely 180 bases long and contain all degenerate nucleotides at every nucleotide position, enabling the introduction of three non-wildtype nucleotides at each position of interest. We divided each exon into specific regions, typically ranging from 30-50 bp in length, to adapt to the availability of sgRNAs and genomic regions of interest. If proximate sgRNAs were not readily available, we utilized a combination of two sgRNAs to cover the intervening region. Additionally, each ssODN was engineered to include a fixed synonymous PAM mutation, preventing subsequent Cas9 cleavage and serving as a consistent HDR marker for later sequencing analysis.

To reduce variability of knock-in efficiency, the same set of sgRNA and oligo library was used for three nucleofections each containing 3.3 ×10^6^ mESCs. A total of 10 x 10^6^ mESCs from these three replicates were pooled into one 10 cm dish after nucleofection, which represents replicate 1. The same experimental strategy was repeated to yield replicate 2. The cells were cultured post-nucleofection, for an initial 72-hour period, with regular media changes every 24 hours. After this, we trypsinized each plate. Half of the cell population was collected for DNA isolation, while the remaining cells were re-plated onto a single 10 cm feeder plate and cultured for an additional 4 days of culture, with daily media changes. On the 7th day following nucleofection, we plated 10^6^ cells from each plate onto three separate 10 cm feeder-containing dishes, preparing them for subsequent drug treatments. On days 8, 10, and 12, we added fresh media supplemented with either 0.4 µM cisplatin or 0.05 µM olaparib, in addition to a DMSO control. Following the drug treatment phase, cells from both replicates for each oligo pool were consolidated on day 14 and pelleted to facilitate DNA extraction.

Genomic DNA was extracted from the cell pellets utilizing the Zymo Genomic DNA Extraction kit (Cat# D3024), and targeted regions were PCR amplified, purified and sent for Next-Generation Sequencing (NGS) at Azenta/Genewiz (New Jersey). The DNA libraries were meticulously prepared in line with the protocol specified in the Illumina TruSeq Nano DNA Library Prep kit and subsequently underwent deep sequencing on the MiSeq platform, employing a 2 x 250 cycle sequencing method.

### 4. Variant analysis and functional classification

Paired-end sequencing was performed on an Illumina MiSeq instrument with a read coverage of 1-3 million reads per sample. The reads were demultiplexed using bcl2fastq (Illumina) and fastq files were generated. SeqPrep was used for adapter trimming and merging the read pairs. Merged reads containing “N” bases were removed from the analysis. The remaining reads were aligned to GRCh38 human reference genome using the needleall command from the EMBOSS package. Abundances of SNVs were quantified when the reads contained the synonymous PAM modification (HDR marker) and no other substitution or indels were present. The variants generated at a low frequency (less than 1 in 10^5^ reads in any one of the replicates) were excluded from further analysis to prevent erroneous variant classification. A pseudocount of 1 was added to all reads, for every sample and at all the conditions. Read counts for each SNVs were then normalized to the total read coverage of the sequencing library. Individual variants with a read count of more than 1 in 100,000 reads at day 3 were used for analysis. The dropout or enrichment scores were calculated by taking the ratio of frequency of the variants at day 14 in DMSO or cisplatin or olaparib over day 3. The scores were expressed in log_2_ scale, which we define as function scores of SNVs in DMSO, cisplatin and olaparib.

### 5. Statistical modeling for SNV classification by AVENGERS

The ratios of frequency of each SNV at day 14 over day 3 were normalized within each experiment using the median ratio for synonymous and non-sense SNV. Position biases in editing rates were modelled using these ratios and the chromosomal position of each SNV within each exon. The ratio was expressed as a function of chromosomal position using the “loess” function from R. The score obtained from the modeling was then subtracted from the log2 ratio, thus removing any positional bias. The corrected log2 ratios were linearly normalized within each exon and across all exons, such that the median nonsense function scores within an exon matched the global nonsense function scores across all exons. For each SNV, four function scores were calculated: one for each treatment (DMSO, cisplatin, olaparib) and one global using a weighted mean of the three conditions (weights: 0.4 for DMSO, 0.25 for cisplatin and olaparib).

The position-corrected and normalized function scores were used as an input for a Gaussian mixture model estimating the probability for each SNV to be functional or non-functional. The synonymous variants that were not expected to affect splicing (based on SpliceAI predictions) constituted the “Functional” class and the non-sense variants constituted the “Non-functional" class. A model was trained using this dataset and the “mclust” function in R. Then, the model predicted the probability of pathogenicity for all SNV using theirs function scores. SNV with a probability of pathogenicity higher than 95% were considered “Non-functional” and SNV with a probability of pathogenicity lower than 1% were considered “Functional”. The remaining SNV were considered as “Uncertain”. The classification obtained using DMSO data were considered for final classification. The variants that showed discordance between DMSO and global (weighted mean) classifications were classified as “Uncertain”.

ROC curves were used to assess performance of the computational model at predicting assigned ClinVar classifications using SGE data and other predictors (CADD^44^, BayesDel^45^, REVEL^46^, PRIOR, EVE^47^ and AlphaMissense^48^). ROC curves were produced with the package “pROC” in R and using missense pathogenic and benign variants as predictors. The SGE function scores were also used to predict “non-sense” and “synonymous” codon types.

### 6. Saturation scan using MAVISp for variant interpretation

We applied the simple mode of the MAVISp framework^49^ to retrieve and aggregate missense mutations in COSMIC^50^, cBioPortal^51^, and ClinVar^36^ for BRCA2 using the Uniprot identifier P51587 and the RefSeq identifier NP_000050. Within the MAVISp framework, we used free energy calculations to predict the effects of the variants on folding/unfolding or binding free energies^57,58^. In addition, folding free energy data for all potential mutations within BRCA2 region spanning residues 2470-3185 were obtained using MutateX and RosettaDDGP prediction with the cartesian2020 protocol and ref2015 energy function^57,58^. We classified the mutations for changes in folding or binding free energy according to a consensus approach between the two protocols applied for the free energy calculations, as previously described^49^. For the calculations, we used the AlphaFold model of BRCA2 (residues 2479-3185) from the AlphaFold database^59^. The structure of the complex between BRCA2 and SEM1 has been modelled with AF-multimer by using the murine BRCA2 C-terminal domain 2479-3185 and the whole sequence of SEM1 (i.e., residues 1-70). The quality of the model has been assessed through superimposition with the structure of the C-terminal domain of BRCA2 and SEM1 contained in the PDB entry 1IYJ^60^. We also retrieved or calculated the pathogenicity scores from AlphaMissense^48^, DeMask^48^, and EVE^47^ using the tools implemented in MAVISp. The final classification of each amino acid in the HD, OB1, OB2 and OB3 domains were mapped along with AlphaMissense and EVE classifications in the form of a heatmap presenting the classification for each amino acid in these domains, using a custom R script.

### 7. Function scores mapping to the BRCA2 PDB structure

Function scores for all SNVs were mapped onto the structures of *BRCA2* by averaging missense SNV probability of pathogenicity at each amino acid position. The amino acids of the PDB structure: 1MIU was aligned with the corresponding amino acids in human BRCA2 sequence using COBALT. For each amino acid, the averaged probability of pathogenicity was calculated using the missense SNV affecting the given amino acid. The figure was then created using Jmol and POV-Ray.

## Author contribution

SS and SKS conceived the study and designed the experimental approach, MG performed the bioinformatic analysis, SS, ES, DC, TLS performed the experiments, JG performed cloning of sgRNA, MA and EP performed MAVISp analyses, RC helped in genome editing experiments, SS and SKS wrote the manuscript, and all authors were involved during the discussion and editing.

## Acknowledgements

We thank Drs. Kajal Biswas, Rishabh Sharan, Sambita Modak, Debojyoti Chakraborty, Sundaram Acharya and members of our laboratory for helpful discussions and suggestions on this project. We acknowledge the support from Jeff Carrel and Megan Karwan from the CCR flow cytometry core, Elizabeth Conner from CCR Genomics core. Figures in this manuscript are made using Adobe Illustrator v6 and illustrations were made using a paid subscription to BioRender.com.

## Conflict of interest

The authors declare no competing or financial interests.

## Funding

This project has been funded by the Intramural Research Program, Center for Cancer Research, National Cancer Institute, U.S. National Institutes of Health (SKS). The EP group is supported by Danmarks Grundforskningsfond (DNRF125), and the MAVISp calculations have been supported by a EuroHPC Benchmark Access Grant (EHPC-BEN-2023B02-010) on Discoverer. MA is supported by a PhD Fellowship from the Danish Data Science Academy (DDSA).

## Notes

### Competing Interest Statement

The authors have declared no competing interest.

## References

1. Claussnitzer, M. et al. A brief history of human disease genetics. Nature 577, 179–189 (2020).

2. Dorling, L. et al. Breast cancer risks associated with missense variants in breast cancer susceptibility genes. Genome Med. 14, 51 (2022).

3. Breast Cancer Association Consortium et al. Breast Cancer Risk Genes - Association Analysis in More than 113,000 Women. N. Engl. J. Med. 384, 428–439 (2021).

4. Nik-Zainal, S. et al. Landscape of somatic mutations in 560 breast cancer whole-genome sequences. Nature 534, 47–54 (2016).

5. Cooper, G. M. & Shendure, J. Needles in stacks of needles: finding disease-causal variants in a wealth of genomic data. Nat. Rev. Genet. 12, 628–40 (2011).

6. Tabet, D., Parikh, V., Mali, P., Roth, F. P. & Claussnitzer, M. Scalable Functional Assays for the Interpretation of Human Genetic Variation. Annu. Rev. Genet. 56, 19.1–19.25 (2022).

7. Starita, L. M. et al. Variant Interpretation: Functional Assays to the Rescue. Am. J. Hum. Genet. 101, 315–325 (2017).

8. Weile, J. & Roth, F. P. Multiplexed assays of variant effects contribute to a growing genotype-phenotype atlas. Hum. Genet. 137, 665–678 (2018).

9. Fowler, D. M. & Fields, S. Deep mutational scanning: a new style of protein science. Nat. Methods 11, 801–7 (2014).

10. Findlay, G. M. et al. Accurate classification of BRCA1 variants with saturation genome editing. Nature 562, 217–222 (2018).

11. Findlay, G. M., Boyle, E. A., Hause, R. J., Klein, J. C. & Shendure, J. Saturation editing of genomic regions by multiplex homology-directed repair. Nature 513, 120–3 (2014).

12. Hanna, R., et al. Massively parallel assessment of human variants with base editor screens. Cell 184, 1064-1080.e20 (2020).

13. Cuella-Martin, R. et al. Functional interrogation of DNA damage response variants with base editing screens. Cell 184, 1081–1097.e19 (2021).

14. Huang, C., Li, G., Wu, J., Liang, J. & Wang, X. Identification of pathogenic variants in cancer genes using base editing screens with editing efficiency correction. Genome Biol. 22, 80 (2021).

15. Erwood, S. et al. Saturation variant interpretation using CRISPR prime editing. Nat. Biotechnol. 40, 885–895 (2022).

16. Monteiro, A. N. et al. Variants of uncertain clinical significance in hereditary breast and ovarian cancer genes: best practices in functional analysis for clinical annotation. J. Med. Genet. jmedgenet-2019-106368 (2020) doi:10.1136/jmedgenet-2019-106368.

17. Parsons, M. T. et al. Large scale multifactorial likelihood quantitative analysis of BRCA1 and BRCA2 variants: An ENIGMA resource to support clinical variant classification. Hum. Mutat. 40, 1557–1578 (2019).

18. Guidugli, L. et al. Assessment of the Clinical Relevance of BRCA2 Missense Variants by Functional and Computational Approaches. Am. J. Hum. Genet. 102, 233–248 (2018).

19. Richardson, M. E. et al. Strong functional data for pathogenicity or neutrality classify BRCA2 DNA-binding-domain variants of uncertain significance. Am. J. Hum. Genet. 108, 458–468 (2021).

20. Biswas, K. et al. A computational model for classification of BRCA2 variants using mouse embryonic stem cell-based functional assays. NPJ genomic Med. 5, 52 (2020).

21. Ikegami, M. et al. High-throughput functional evaluation of BRCA2 variants of unknown significance. Nat. Commun. 11, 2573 (2020).

22. Biswas, K. et al. Sequencing-based functional assays for classification of BRCA2 variants in mouse ESCs. Cell Reports Methods 100628 (2023) doi:10.1016/j.crmeth.2023.100628.

23. Mishra, A. P. et al. BRCA2-DSS1 interaction is dispensable for RAD51 recruitment at replication-induced and meiotic DNA double strand breaks. Nat. Commun. 13, 1751 (2022).

24. Mishra, A. P. et al. Characterization of BRCA2 R3052Q variant in mice supports its functional impact as a low-risk variant. Cell Death Dis. 14, 753 (2023).

25. Hartford, S. A. et al. Interaction with PALB2 Is Essential for Maintenance of Genomic Integrity by BRCA2. PLoS Genet. 12, e1006236 (2016).

26. von Nicolai, C., Ehlén, Å., Martin, C., Zhang, X. & Carreira, A. A second DNA binding site in human BRCA2 promotes homologous recombination. Nat. Commun. 7, 12813 (2016).

27. Yang, H. et al. BRCA2 function in DNA binding and recombination from a BRCA2-DSS1-ssDNA structure. Science 297, 1837–48 (2002).

28. Petrucelli, N., Daly, M. B. & Pal, T. BRCA1- and BRCA2-Associated Hereditary Breast and Ovarian Cancer. GeneReviews® (1993).

29. Kuchenbaecker, K. B. et al. Risks of Breast, Ovarian, and Contralateral Breast Cancer for BRCA1 and BRCA2 Mutation Carriers. JAMA 317, 2402–2416 (2017).

30. Roy, R., Chun, J. & Powell, S. N. BRCA1 and BRCA2: different roles in a common pathway of genome protection. Nat. Rev. Cancer 12, 68–78 (2011).

31. Jensen, R. B., Carreira, A. & Kowalczykowski, S. C. Purified human BRCA2 stimulates RAD51-mediated recombination. Nature 467, 678–83 (2010).

32. Esashi, F. et al. CDK-dependent phosphorylation of BRCA2 as a regulatory mechanism for recombinational repair. Nature 434, 598–604 (2005).

33. Kuznetsov, S. G., Liu, P. & Sharan, S. K. Mouse embryonic stem cell-based functional assay to evaluate mutations in BRCA2. Nat. Med. 14, 875–81 (2008).

34. Sharan, S. K. BRCA2 deficiency in mice leads to meiotic impairment and infertility. Development 131, 131–142 (2003).

35. Sahu, S. et al. Saturation genome editing of 11 codons and exon 13 of BRCA2 coupled with chemotherapeutic drug response accurately determines pathogenicity of variants. PLoS Genet. 19, e1010940 (2023).

36. Landrum, M. J. et al. ClinVar: public archive of relationships among sequence variation and human phenotype. Nucleic Acids Res. 42, D980–5 (2014).

37. Cline, M. S. et al. BRCA Challenge: BRCA Exchange as a global resource for variants in BRCA1 and BRCA2. PLoS Genet. 14, e1007752 (2018).

38. Lord, C. J. & Ashworth, A. PARP inhibitors: Synthetic lethality in the clinic. Science 355, 1152–1158 (2017).

39. Richards, S. et al. Standards and guidelines for the interpretation of sequence variants: a joint consensus recommendation of the American College of Medical Genetics and Genomics and the Association for Molecular Pathology. Genet. Med. 17, 405–24 (2015).

40. Brnich, S. E. et al. Recommendations for application of the functional evidence PS3/BS3 criterion using the ACMG/AMP sequence variant interpretation framework. Genome Med. 12, 3 (2019).

41. Sahu, S. et al. Protocol for the saturation and multiplexing of genetic variants using CRISPR-Cas9. STAR Protoc. 4, 102702 (2023).

42. Sirisena, N. et al. Functional evaluation of five BRCA2 unclassified variants identified in a Sri Lankan cohort with inherited cancer syndromes using a mouse embryonic stem cell-based assay. Breast Cancer Res. 22, 43 (2020).

43. Biswas, K. et al. A comprehensive functional characterization of BRCA2 variants associated with Fanconi anemia using mouse ES cell-based assay. Blood 118, 2430–42 (2011).

44. Rentzsch, P., Schubach, M., Shendure, J. & Kircher, M. CADD-Splice-improving genome-wide variant effect prediction using deep learning-derived splice scores. Genome Med. 13, 31 (2021).

45. Feng, B.-J. PERCH: A Unified Framework for Disease Gene Prioritization. Hum. Mutat. 38, 243–251 (2017).

46. Ioannidis, N. M. et al. REVEL: An Ensemble Method for Predicting the Pathogenicity of Rare Missense Variants. Am. J. Hum. Genet. 99, 877–885 (2016).

47. Frazer, J. et al. Disease variant prediction with deep generative models of evolutionary data. Nature 599, 91–95 (2021).

48. Cheng, J. et al. Accurate proteome-wide missense variant effect prediction with AlphaMissense. Science 381, eadg7492 (2023).

49. Arnaudi, M., et al. MAVISp: Multi-layered Assessment of VarIants by Structure for proteins. bioRxiv 2022.10.22.513328 (2023) doi:10.1101/2022.10.22.513328.

50. Tate, J. G. et al. COSMIC: the Catalogue Of Somatic Mutations In Cancer. Nucleic Acids Res. 47, D941–D947 (2019).

51. de Bruijn, I. et al. Analysis and Visualization of Longitudinal Genomic and Clinical Data from the AACR Project GENIE Biopharma Collaborative in cBioPortal. Cancer Res. (2023) doi:10.1158/0008-5472.CAN-23-0816.

52. Mishra, A. P. et al. BRCA2-DSS1 interaction is dispensable for RAD51 recruitment at replication-induced and meiotic DNA double strand breaks. Nat. Commun. 13, 1751 (2022).

53. Guo, Q. et al. Functional evaluation of BRCA1/2 variants of unknown significance with homologous recombination assay and integrative in silico prediction model. J. Hum. Genet. (2023) doi:10.1038/s10038-023-01194-6.

54. Hart, S. N. et al. Comprehensive annotation of BRCA1 and BRCA2 missense variants by functionally validated sequence-based computational prediction models. Genet. Med. 21, 71–80 (2019).

55. Shimelis, H. et al. BRCA2 Hypomorphic Missense Variants Confer Moderate Risks of Breast Cancer. Cancer Res. 77, 2789–2799 (2017).

56. Ran, F. A. et al. Genome engineering using the CRISPR-Cas9 system. Nat. Protoc. 8, 2281–2308 (2013).

57. Tiberti, M. et al. MutateX: an automated pipeline for in silico saturation mutagenesis of protein structures and structural ensembles. Brief. Bioinform. 23, (2022).

58. Sora, V. et al. RosettaDDGPrediction for high-throughput mutational scans: From stability to binding. Protein Sci. 32, e4527 (2023).

59. Jumper, J. et al. Highly accurate protein structure prediction with AlphaFold. Nature 596, 583–589 (2021).

60. Yang, H. et al. BRCA2 Function in DNA Binding and Recombination from a BRCA2-DSS1-ssDNA Structure. Science (80-.). 297, 1837–1848 (2002).

61. Walker, L. C. et al. Using the ACMG/AMP framework to capture evidence related to predicted and observed impact on splicing: Recommendations from the ClinGen SVI Splicing Subgroup. Am. J. Hum. Genet. 110, 1046–1067 (2023).

