## Supplementary Figures 1-9 for "AVENGERS: Analysis of Variant Effects using Next Generation sequencing to Enhance *BRCA2* Stratification"

**Supplementary figure 1: Generation of all possible SNVs using CRISPR-Cas9 based SGE.**

**(a)** Heatmap showing the total number of expected and recovered variants across 48 experimental pools to saturate the C-terminal DNA binding domain. The legend represents the number of variants ranging from 87 SNVs (light blue) to 339 SNVs (dark blue). A total of 95.8% of all possible SNVs were recovered (6270 out of 6544 SNVs). **(b-c)** Bar plot showing the distribution of (b) indel and (c) HDR rates calculated for each experimental pool. HDR rates are calculated based on the percentage of reads for SNVs with fixed PAM modification in each experimental pool. (d) Pearson correlation between the read counts of all SNVs recovered after HDR at a frequency of 1 in $10^5$ reads in both replicates across the experimental pools.

Supplementary figure-1

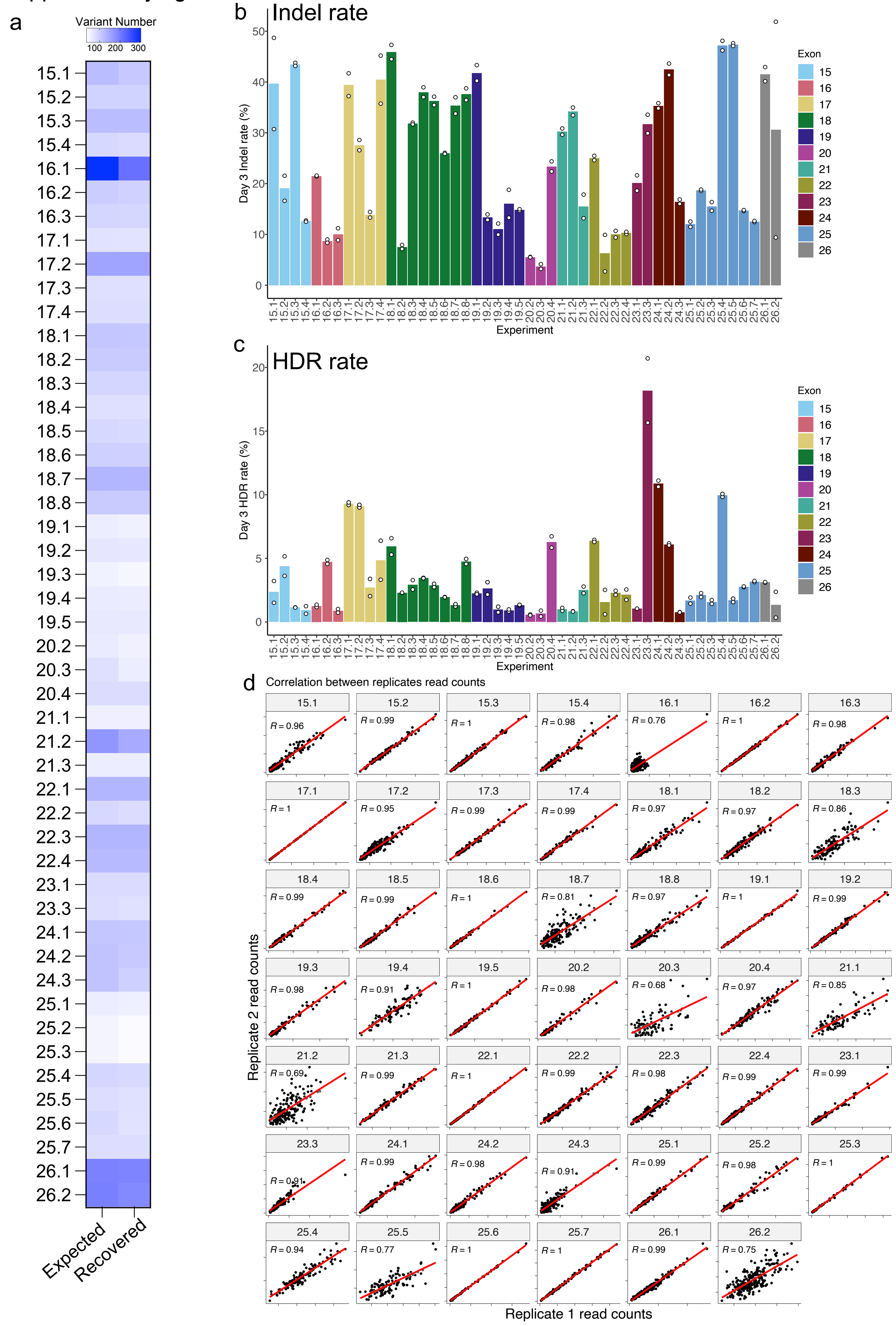

**Supplementary figure 2: Function scores based on cell fitness and drug response enhances classification accuracy of AVENGERS.**

**(a)** The Pearson correlation of Function Scores (FS) for 6270 SNVs between cell fitness (DMSO treated) and response to drugs (cisplatin treated). Additionally, we assessed the correlation between cell fitness and olaparib treatment, as well as the correlation between cisplatin and olaparib responses. **(b)** Schematic showing our filtering strategy to integrate classification scores based on cell fitness and drug response data to accurately categorize SNVs. 6270 SNVs were categorized using Mixture-modelling based on their survival at day 14, and integrated FS values were computed by combining cell fitness and drug response scores. **(c)** Gaussian Mixture Modeling revealed that 5968 SNVs (95.1%, 4881 functional and 1087 non-functional) concurred with cell fitness data. Only 302 SNVs (4.8%) had conflicting classifications, so we excluded them as "uncertain" **(d)** Heatmap showing the correlation matrix of the read counts and the FS for each experimental pool. The legend represents the correlation value ranging from 0.15 (white) to 1 (red). The second heatmap represents the percentage of variant recovered and then finally classified by integrating the scores and excluding the uncertain variants. The legend represents the percentage ranging from 83% (light blue) to 100% (dark blue).

Supplementary figure-2

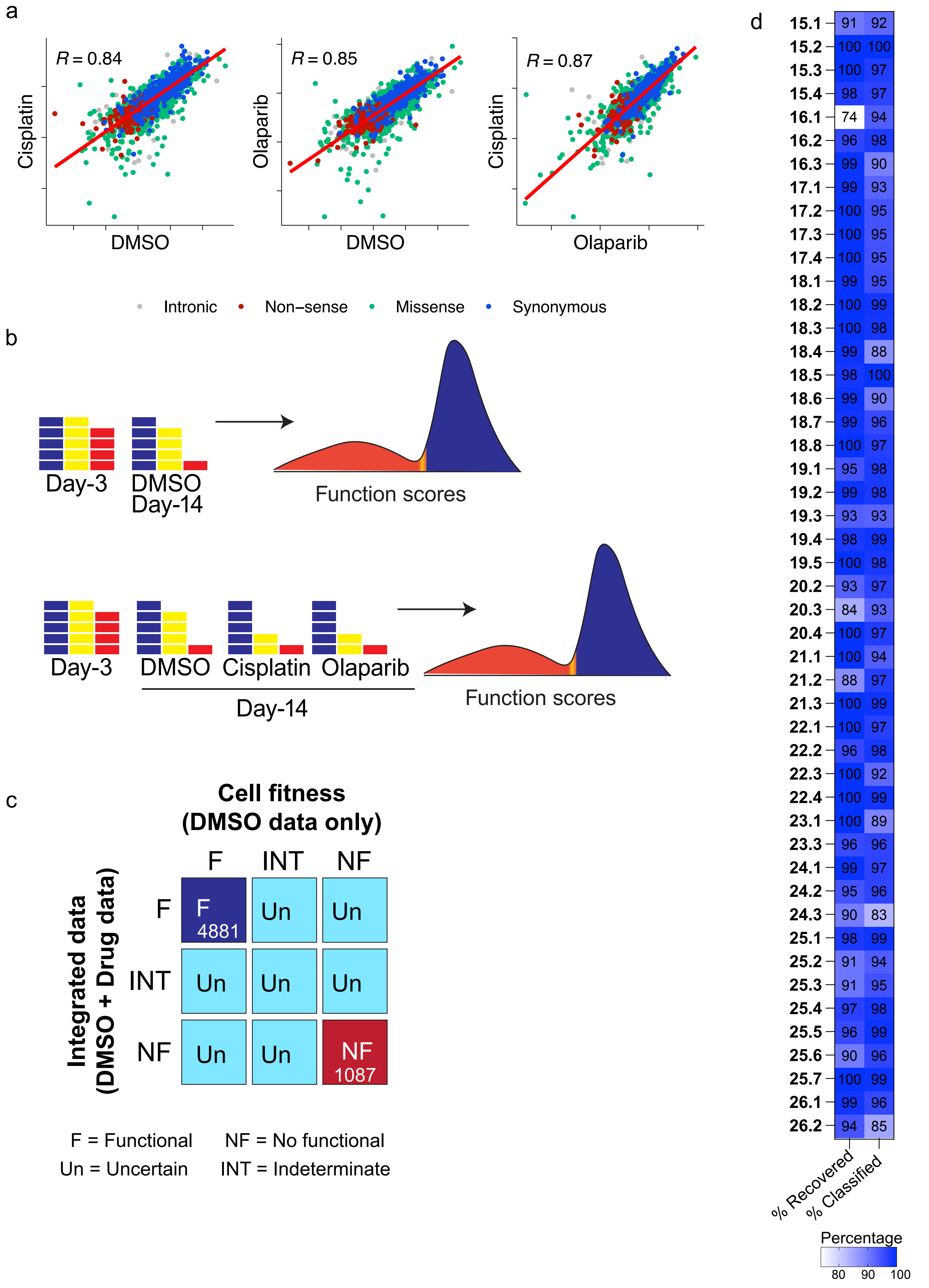

**Supplementary figure 3: SGE experiments reveal temporal depletion of Cas9-induced indels.**

Histogram showing the distribution of indels identified in each experimental pool, along with their corresponding function scores. In this analysis, we considered indels that met specific criteria: they had to align with the reference genome, displaying a single insertion or deletion within 50 base pairs of the anticipated Cas9 cleavage site defined by the sgRNAs used in each experimental pool. These indels were only counted if their frequency was at least 1 in $10^6$ reads in both replicates, and in-frame indels were marked when their size was divisible by 3. Function scores were determined by comparing the frequency of indels at day 14 relative to day 3. We observed depletion of frameshifting indels in the pool, while a few in-frame indels were occasionally tolerated.

Supplementary figure-3

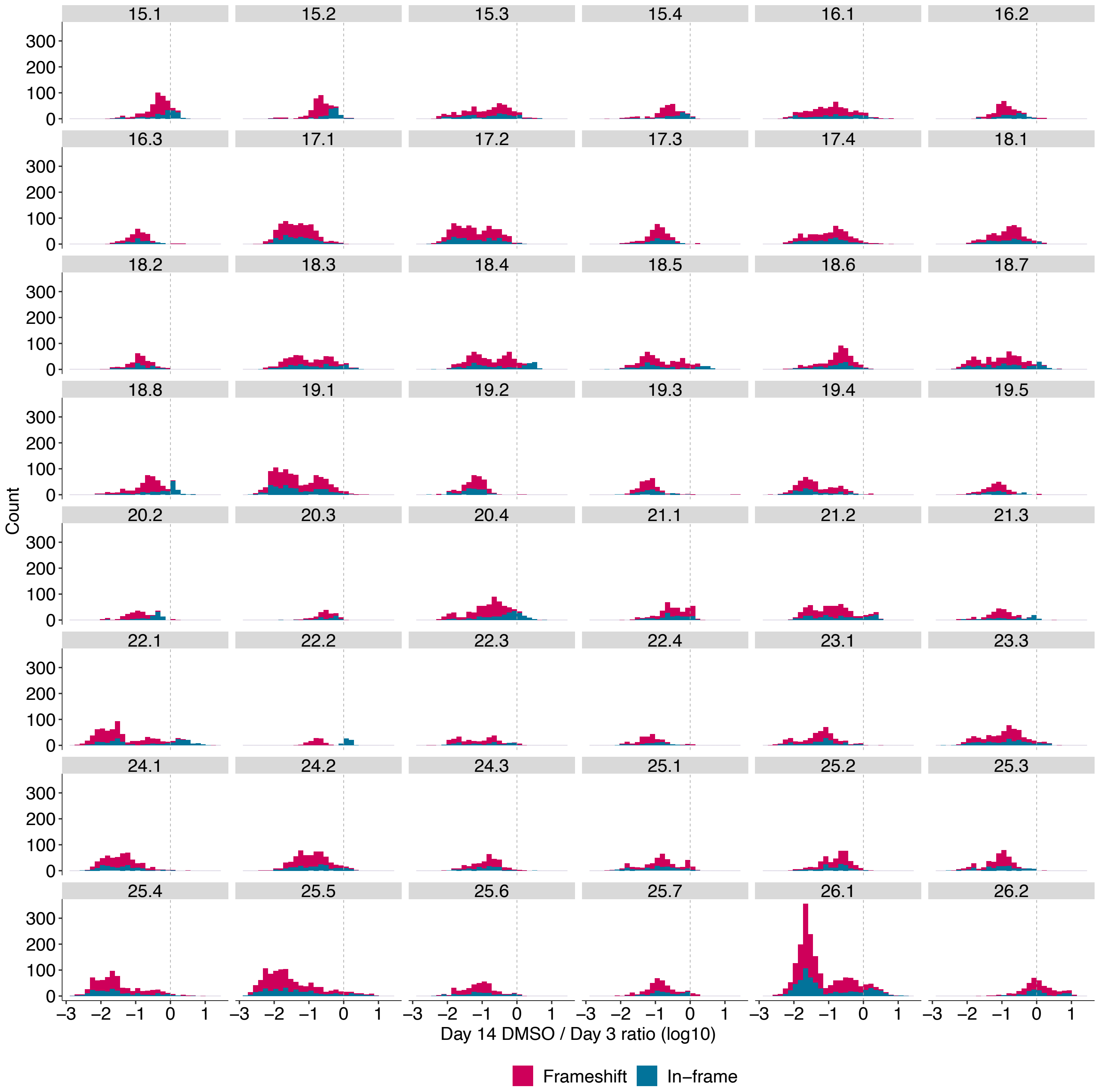

**Supplementary figure 4: Mixture modelling of function scores to classify *BRCA2* SNVs.**

**(a)** Gaussian Mixture modelling (GMM) plot showing the distribution of probability of impact on function (PIF) and SNV function scores categorized between synonymous (blue), nonsense (red), intronic (grey) and missense SNVs across CTD. **(b)** Correlation of function scores of all SNVs between individual replicates demonstrate the distribution of functional and nonfunctional thresholds. **(c)** Correlation of function scores of all SNVs between individual replicates for individual exons. The dashed line represents the thresholds for functional and nonfunctional thresholds derived from GMM. **(d-f)** Swarm plots demonstrating the distribution of (d) synonymous and nonsense SNVs (n = 1473) (e) ClinVar-reported SNVs (n = 696) and (f) all classified variants (n = 5968). The dashed line represents the median of the nonsense SNVs (red) and synonymous SNVs (blue).

### Supplementary figure-4

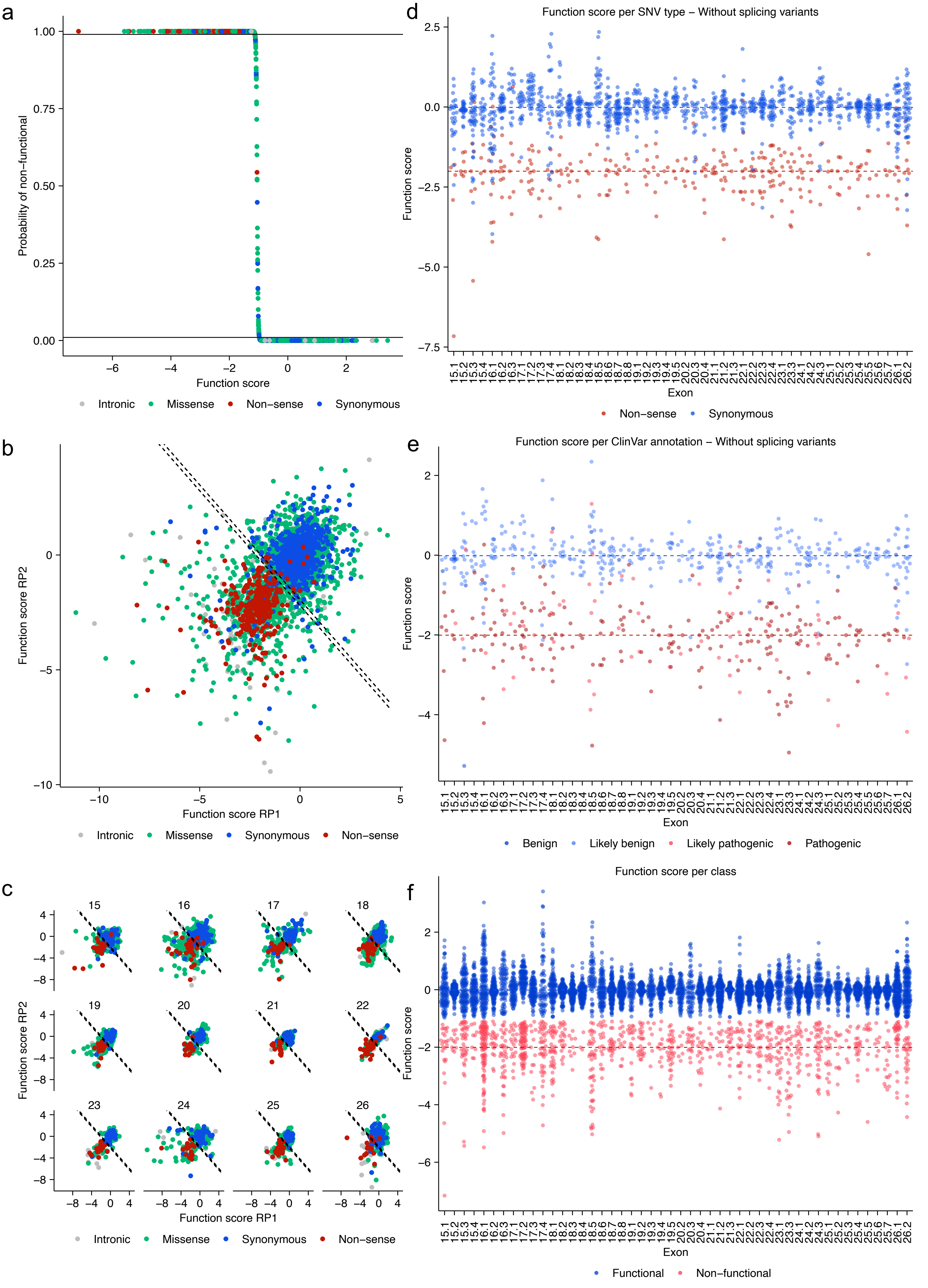

**Supplementary figure 5: AVENGERS outperforms computational meta predictors. (a)** ROC curves indicate the performance of computational models at categorizing ClinVar-reported missense variants in the *BRCA2* CTDB domain. ROC shows an AUC value of 0.96 in classifying ClinVar variants and AUC value of 0.97 in accurately categorizing nonsense and synonymous SNVS. We observed a moderate concordance to other computational predictors evaluated like CADD[44], BayesDel[45], REVEL[46], PRIOR, EVE[47] and AlphaMissense[48]. **(b)** Correlation between SGE-derived function scores and computational metrics in determining ClinVar-reported BRCA2 SNVs. The color code represents the benign, likely benign, pathogenic, and likely pathogenic class reported in ClinVar (n = 5968 SNVs)

### Supplementary figure-5

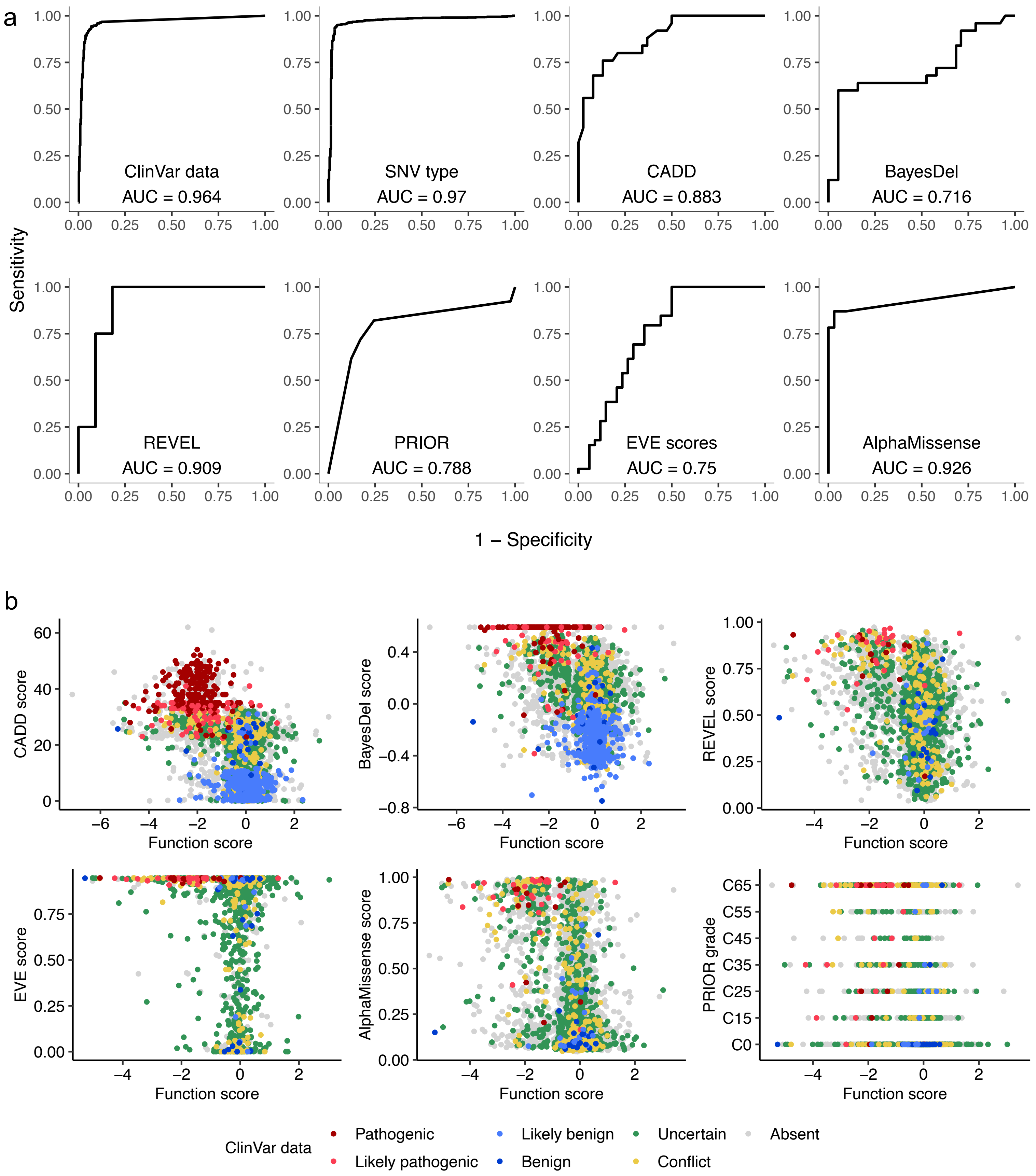

**Supplementary figure 6: AVENGERS map for key functional domains reveal strong concordance to AlphaMissense prediction.**

Sequence-function map displaying the concordance of functional and non-functional categorization of individual amino acid changes across **(a)** helical domain residues 2479-2668, **(b)** OB1 residues from 2682-2794, **(c)** OB2 residues from 2804-3054 and **(d)** OB3 domain residues from 3073-3167 of the BRCA2 CTDB domain. The first box represents AVENGERS dataset and their concordance with MAVISP saturation scan and Alpha-Missense prediction. The box color signifies "functional (blue)", "non-functional (red)", and "uncertain (turquoise blue)" class. The wildtype amino acid is shown as a circle, while white boxes indicate excluded amino acids. The Y-axis denotes the alternative amino acid changes. The magnified image of the 3D ribbon plot depicts the position of amino acid residues where clusters of functional/non-functional SNVs are reported across individual domains.

Supplementary figure-6

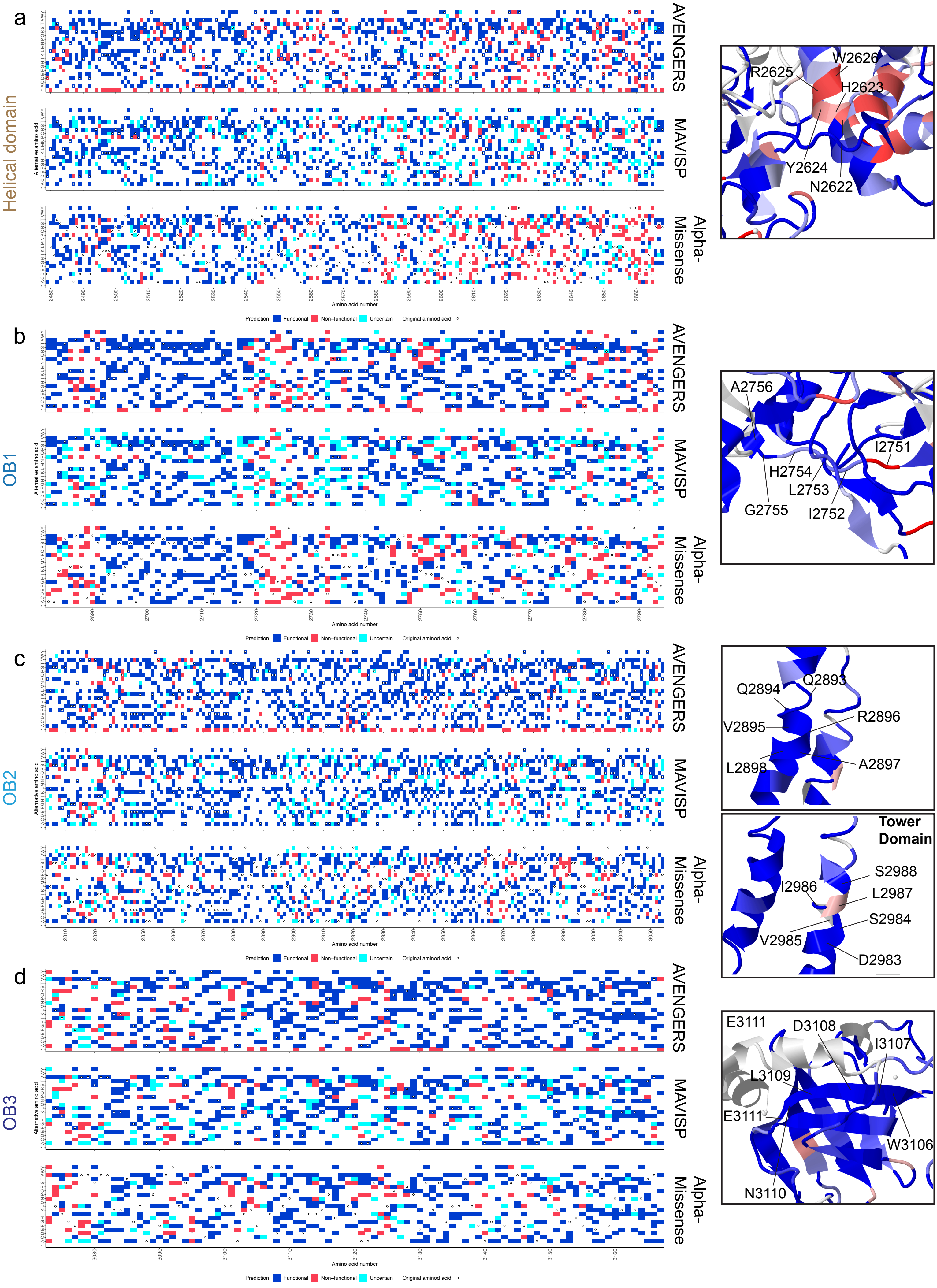

**Supplementary figure 7: AVENGERS is concordant with orthogonal functional assay.**

**(a)** Strong concordance of CRISPR-based SGE classification with CRISPR-based Prime Editing (n=156 SNVs). **(b)** Comparison of SGE classification with CRISPR-independent methods such as BAC recombineering and cDNA-based functional classification. Notably, 91.6% of the SNV classifications in our SGE dataset, which includes 75 functional and 56 nonfunctional variants, were in agreement with BAC recombineering results. **(c)** Comparison of SGE classification for 117 SNVs previously classified using the MANO-B assay based on their responses to various PARP inhibitors (olaparib, rucaparib, niraparib) and carboplatin. **(d)** Comparison with HDR-based functional classification revealed that 43 out of 47 SNVs matched with our SGE classification. The box color key denotes functional variants in light blue, non-functional in red, and uncertain SNVs in light red.

### Supplementary figure-7

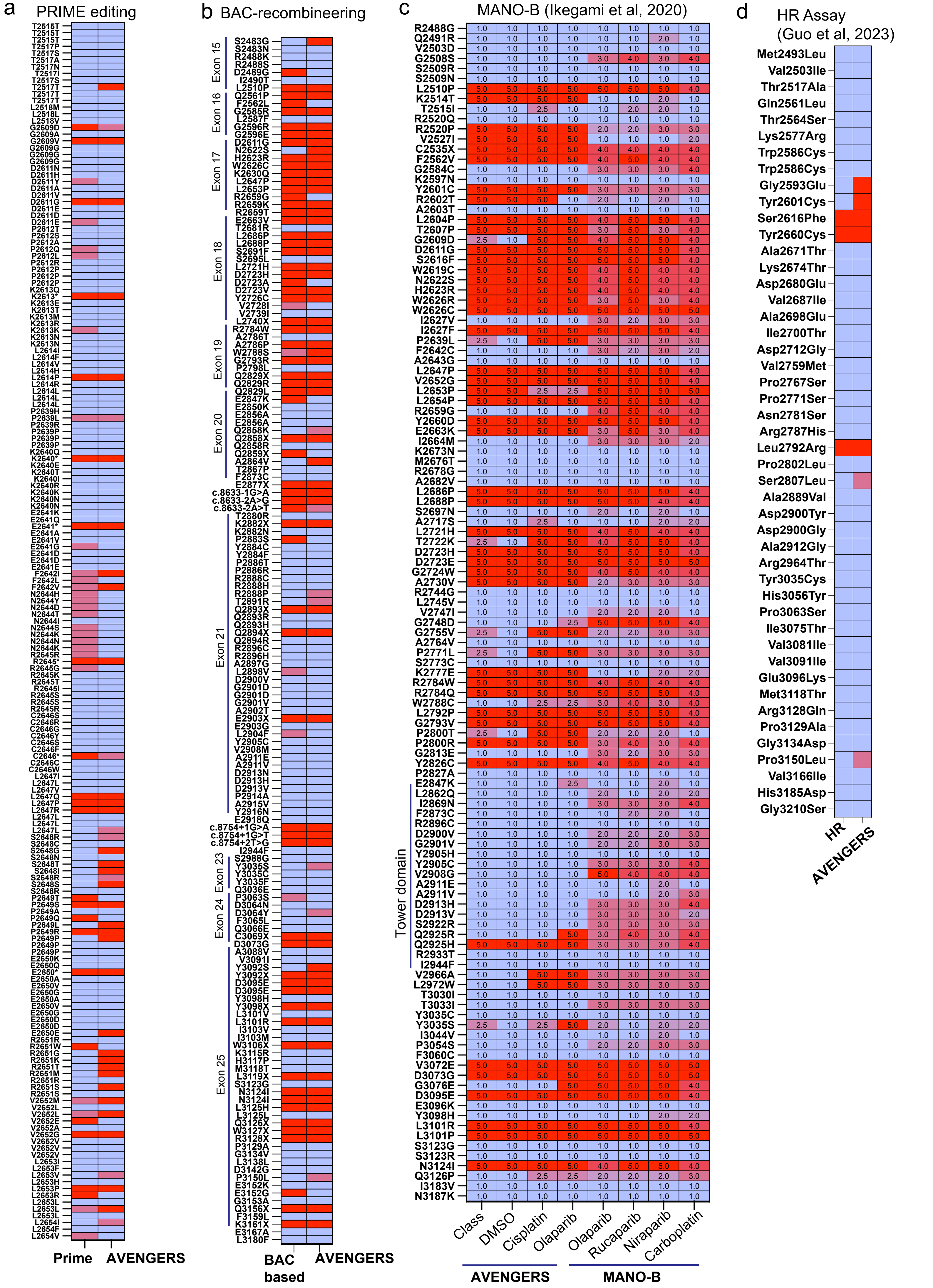

**Supplementary figure 8: SGE functional classification correlates with SNVs ability to perform homologous recombination (HR)**

Robust alignment between the SGE-derived functional classification and the HR score from Richardson et al. 2021, particularly for SNVs distributed across crucial functional regions within the BRCA2 C-terminal DNA binding domain. Notably, SNVs in the Tower domain exhibit a high HR score, consistent with the functional nature of our SGE-based scores (n = 233 SNVs). The box color represents light blue as functional, red as non-functional, and light red as the uncertain class of SNVs.

Supplementary figure-8

a

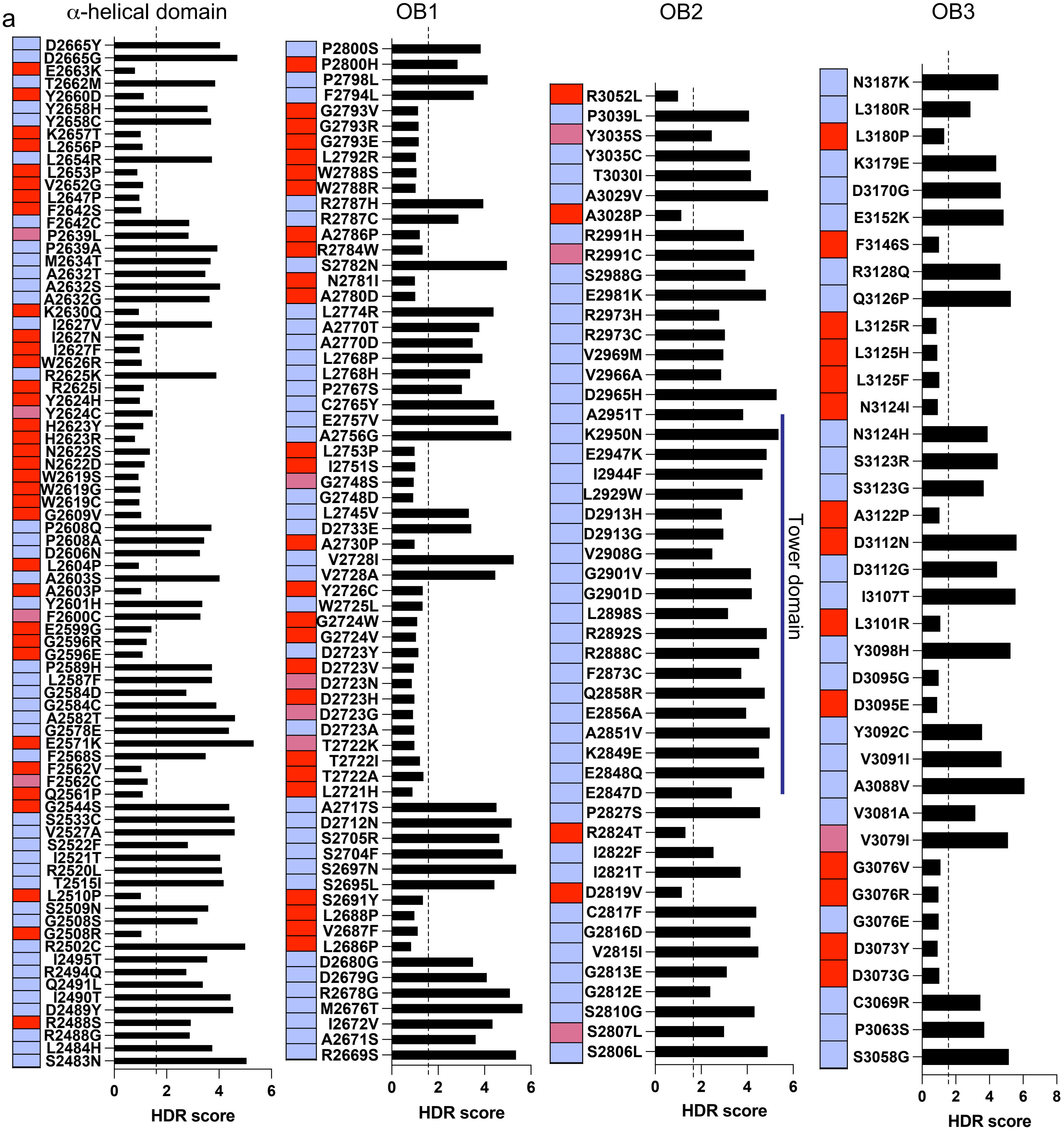

**Supplementary figure 9: Categorizing 149 uncertain SNVs as potential hypomorphic variants.**

**(a)** Experimental strategy demonstrating the filtering of “uncertain” class SNVs with conflicting classifications between cell fitness and drug response data to identify potential hypomorphs (n = 302 SNVs). The uncertain variants were filtered to identify SNVs that fall in the indeterminate and nonfunctional classes and further identified if they represent non-functional class in both cisplatin and olaparib dataset. **(b)** Heatmap showing potential hypomorphic variants that survive in the pool but are sensitive to DNA damaging agents (n = 149 SNVs).

Supplementary figure-9

a

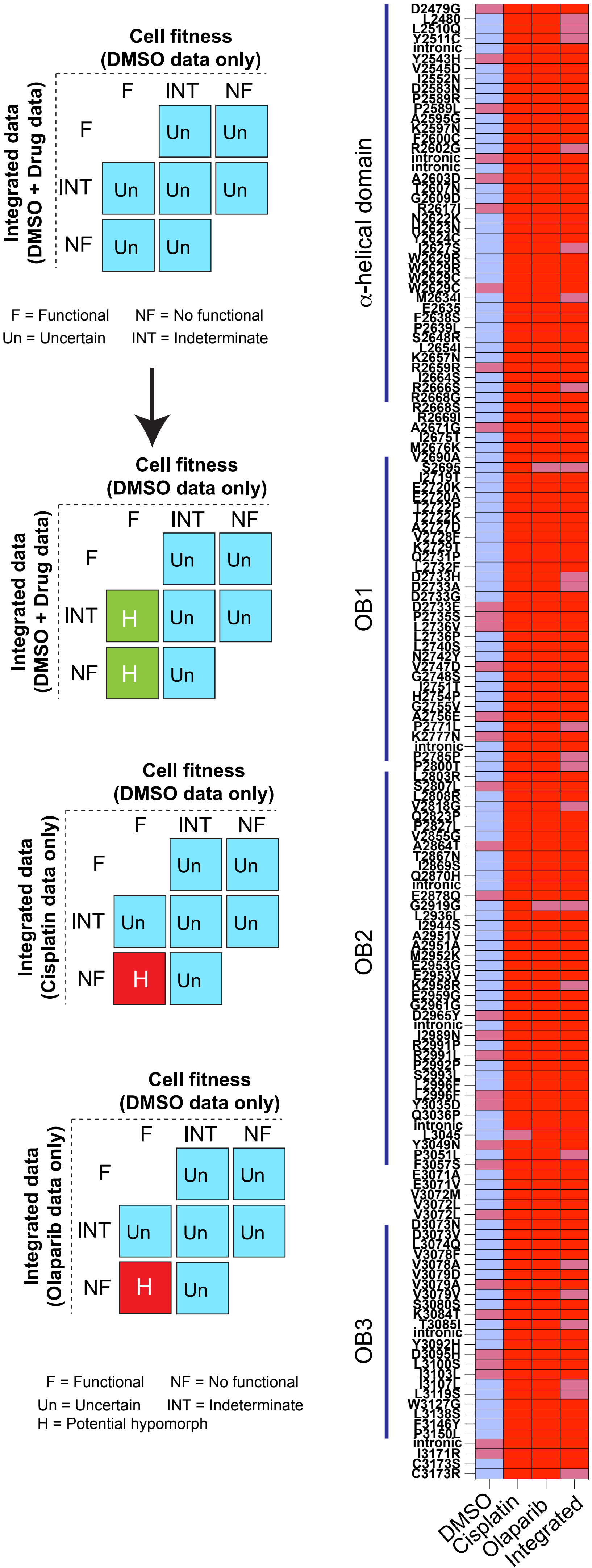
